# INSIDER: Interpretable Sparse Matrix Decomposition for Bulk RNA Expression Data Analysis

**DOI:** 10.1101/2022.11.10.515904

**Authors:** Kai Zhao, Sen Huang, Cuichan Lin, Pak Chung Sham, Hon-Cheong So, Zhixiang Lin

## Abstract

RNA-Seq is widely used to capture transcriptome dynamics across tissues from different biological entities even across biological conditions, with the aim of understanding the contribution of gene activities to phenotypes of biosamples. However, due to variation from tissues and biological entities (or other biological conditions), joint analysis of bulk RNA expression profiles across multiple tissues from a number of biological entities to achieve the aim is hindered. Moreover, it is crucial to consider interactions between biological variables. For example, different brain disorders may affect brain regions heterogeneously. Thus, modeling the disorder-region interaction can shed light on the heterogeneity. To address these key challenges, we propose a general and flexible statistical framework based on matrix factorization, named INSIDER (https://github.com/kai0511/insider).

INSIDER decomposes variation from different biological variables into a shared low-rank latent space. In particular, it considers interactions between biological variables and introduces the elastic net penalty to induce sparsity, thus facilitating interpretation. In the framework, the biological variables and interaction terms can be defined based on the research questions and study design. Besides, it enables us to compute the ‘adjusted’ expression profiles for biological variables that control variation from other biological variables. Lastly, it allows various downstream analyses, such as clustering donors with donor representations, revealing development trajectory in its application to the BrainSpan data, and uncovering mechanisms underlying variables like phenotype and interactions between biological variables (e.g., phenotypes and tissues).

## Introduction

RNA sequencing (RNA-Seq), which has been developed for more than a decade, uses deep sequencing technology to offer a far more precise measurement of levels of transcripts and their isoforms than other methods [1]. It is now widely used to measure transcriptome levels across different tissues from a number of donors and capture transcriptome dynamics across tissues or biological conditions. For example, the Genotype-Tissue Expression (GTEx) [2] project offers a valuable data resource to study tissue-specific gene expression and regulation. Moreover, there are many data resources related to brain development or psychiatric disorders, including the BrainSpan [3], PsychENCODE [4], and ATP (the Autism Tissue Program) [5]. The primary task of these studies is to link gene activities to phenotypes, especially human disorders, and reveal important biological processes (BPs) that contribute to phenotypes.

However, variation in transcript levels from covariates, such as donor and tissue, makes the task challenging. Let us take studies of autism spectrum disorder (ASD) as examples. Different brain regions may demonstrate wide differences in gene activities in ASD [6]. On the other hand, no individuals with ASD have exactly the same genetic expression profiles [7], which is influenced by individual characteristics such as age, sex, diet, and lifestyle. Thus, joint analyses of gene expression by RNA-Seq of multiple brain regions from several donors are hindered by the variation from donors and tissues.

Traditional dimension reduction approaches, such as principal component analysis (PCA), factor analysis, and matrix factorization, cannot handle directly the variation from covariates, such as tissue and donor, to facilitate further analysis. One study [8] proposed a PCA-based approach with adjustment for confounding variables. Other approaches based on factor analysis, such as RUV [9] and RUV-2 [10], have been proposed to adjust for technical artifacts or confounding variables. An empirical Bayesian approach, ComBat, was developed to adjust for batch effects in microarray expression data [11]. RA3 [12], a reference-guided approach based on Bayesian matrix factorization, was proposed to model biological variation in data from single-cell chromatin accessibility sequencing (scCAS) and reference data. Another study [13] proposed an orthogonal projection correction (OPC) method to correct confounders in the classification setting.

Non-negative matrix factorization (NMF) [14]-[16] has gained popularity in the analysis of genomic data. Traditional NMF cannot be directly utilized to address the challenges of confounding. Variants of NMF proposed for the analysis of bulk RNA-Seq data are rather limited, and most of them aim to solve problems from single-cell RNA-Seq [15], [16]. Notably, NMF and its variants fail to directly model down-regulations of elements in the biological setting [17]. This imposes a hurdle in uncovering down-regulated biological processes (BPs). However, the wide applicability of NMF has been demonstrated in the analysis of genomics data. Common and Specific matrix Factorization (CSMF) based on NMF was proposed to learn common patterns from data of multiple interrelated biological scenarios [18]. Another study [19] developed an approach based on coupling two nonnegative matrix factorizations to cluster cells in two samples.

On the other hand, most tensor decomposition approaches [20]-[22] cannot handle missing data or invoke sparsity in latent representations. In addition, Bayesian tensor decomposition approaches [23], [24] face the issues of prior selection and heavy computational burden in analyzing genomic data from RNA-Seq, especially when optimizing with Markov Chain Monte Carlo methods.

To tackle the above challenges, we propose a general and flexible statistical approach, INSIDER, based on matrix decomposition. INSIDER does not assume non-negative constraints and decomposes variation from covariates in data into a shared low-rank latent space. INSIDER has the following virtues. First, INSIDER is a general and flexible framework, in which covariates and interaction terms can be defined based on the research questions and study design. INSIDER can accommodate different biological variables, such as gender, disease status, donor, and other biological conditions. An important feature is that it can handle multiple confounding variables simultaneously. Second, it considers interactions between covariates. For example, in modeling expression changes for different neuropsychiatric disorders, brain region may interact with the type of disorder to affect gene expression levels.

Conventional methods based on matrix factorization usually ignore interactions between biological factors. Moreover, INSIDER allows us to compute the ‘adjusted’ expression that controls for variation in other confounders or covariates, which will be illustrated below. In addition, it introduces the elastic net penalty, which induces sparsity and facilitates interpretability. Finally, INSIDER facilitates different downstream analyses, including clustering donors with donor representations, temporal dynamic analyses, and uncovering disease mechanisms, such as revealing interactions between covariates on the underlying process affecting gene expression. We demonstrate the potential of INSIDER in three real data studies in the Results section.

## Results

### Overview of INSIDER

We propose a novel computational method called INSIDER (<u>in</u>terpretable sparse matrix <u>de</u>composition for bulk <u>R</u>NA expression data analysis), summarized in Figure 1. INSIDER considers that variation in RNA expression levels originates from several biological variables or covariates, such as donor, tissue, and phenotype (Figure 1A). In INSIDER, the variation is decomposed into a shared latent space of rank *K* by matrix factorization (Figure 1B). The effect of each covariate on expression can be measured by multiplying the corresponding latent representation with the gene representation *V* (Figure 1B). Specifically, the expression level of gene *m* in tissue *h* of donor *i* with phenotype *j, z*_*ijhm*_, is modelled as

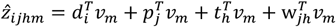

where *d*_*i*_, *p*_*j*_, *t*_*h*_, *w*_*ih*_, *v*_*m*_ are vectors of length *K*, and *w*_*jh*_ is introduced to capture interactions between phenotype *j* and tissue *h*. We assume that variation originated from the covariates and that interactions between the covariates are additive. Let *Z*^*N*×*M*^ denote the observed data matrix for *N* samples and *M* genes. The above equation can be rewritten in the following matrix representation

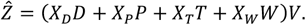

**Figure 1.**
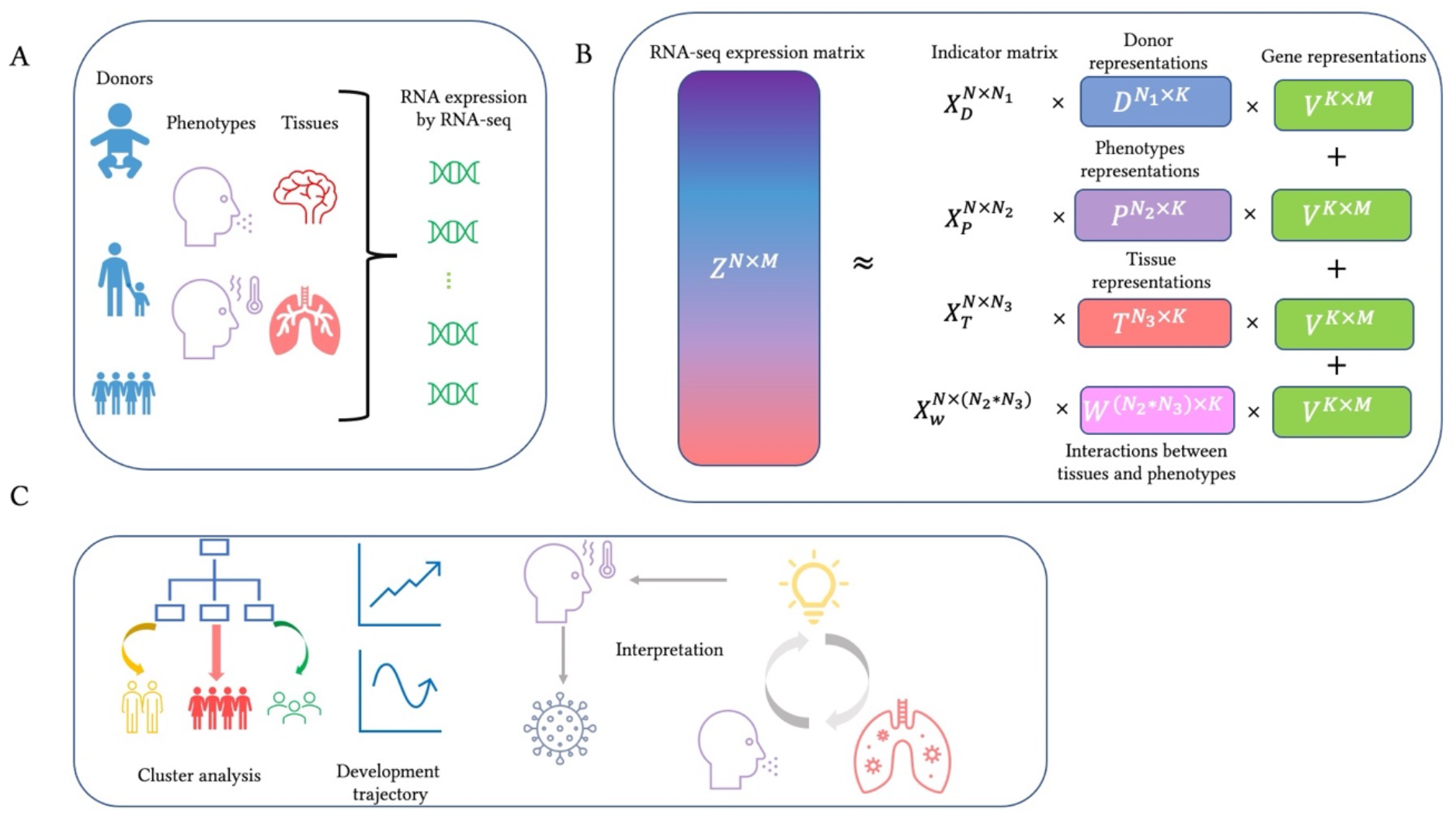
Overview of INSIDER A The RNA-seq data is usually obtained from samples of different tissues from several donors with different phenotypes. B The variation in RNA-seq expression data can be decomposed into a shared *K*-rank latent space by INSIDER. Note that INSIDER incorporates the interaction between covariates and that the gene representation *V* is shared. C Various types of downstream analysis can be conducted on the result from INSIDER. The donor representations can be utilized for cluster analysis, development trajectories can be revealed in the analysis of the BrainSpan data, and biological mechanisms behind phenotypes of interest and interactions between covariates can be revealed. Details on these analyses will be demonstrated in the Result section.

We assume there are *N*_1_ donors, *N*_2_ phenotypes, and *N*_3_ tissues. For the purpose of representation, 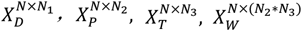 are introduced to be the indicator matrices for donors, phenotypes, tissues, and phenotype by tissue interactions, which represent the dummy design matrices for the samples. 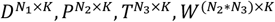 are the matrices for the latent representations for the effects of donors, phenotypes, tissues, and phenotype by tissue interactions on the *K* latent metagenes, respectively, and *V*^*K*×*M*^ represents the effects of *K* metagenes on the expression levels of the *M* genes. Here “metagene” (a common representation) can be conceptualized as certain gene-sets or pathways with heavier weights for some genes than others.

In INSIDER, we seek to minimize the following objective

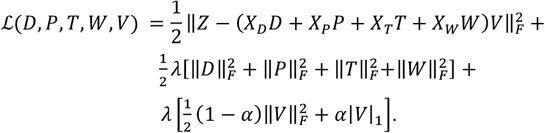

In the above equation, *λ, α* are tuning parameters for the elastic net regularization term, ‖⋅‖_*F*_ represents the Frobenius norm. Alternating block coordinate descent (BCD) is utilized to optimize the objective. INSIDER can incorporate observations with missing values. The technical details of INSIDER are presented in the Methods section.

INSIDER enables various downstream analyses (Figure 1C). Cluster analysis on the donor representation can reveal characteristics of the donors and help find patients subgroups or disease subtyping, development trajectory can be more readily revealed with ‘metagenes’ (low-dimension representations of gene expression) which will be illustrated in its application to the BrainSpan data, and biological processes (BPs) enriched by results from INSIDER help to gain insights into mechanisms underlying covariates like phenotype and interactions between covariates. Details on these analyses are presented below.

### Analysis of the BrainSpan data

The BrainSpan study provides RNA-Seq data profiling up to sixteen cortical and subcortical structures across the entire course of human brain development [25]. In this analysis, we aim to explore the development trajectories across human brain development, the BPs that function in different brain regions, and the involvement of brain regions in different development stages.

The brain span dataset was downloaded from the website provided in [3]. The dimension of the data matrix for our analysis is 524×43411. In this analysis, we modeled development stages and brain regions as covariates but omitted interactions between development stages and brain regions, because many combinations of development stages and brain regions only have 1 or 0 RNA-seq sample, so inference of the interaction will be unreliable. For the definition of development stages, we follow the specifications defined in the original study [3].

For illustration, we denote 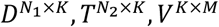 as the latent representations for development stages, brain regions, and genes, respectively. There are *N*_1_ development stages and *N*_2_ brain regions, and *M* is the number of genes included. The rank *K* of latent space selected by hyperparameter tuning is 19.

### Metagenes reveal development trajectories

We adopt the convention to consider the latent representations as metagenes. We first looked at 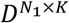, the representation for development stages. We chose the most variable (16^th^), 2^nd,^ and 17^th^ metagenes as our study objects by calculating variance for each column of the matrix *D*. We visualized the trajectories of the selected metagenes across human brain development (Figure 2A). The expression level of the most variable metagene is high before late fetal stages and decreases to a low level after early childhood (Figure 2A). Meanwhile, we see a rapid surge in the level of the 2^nd^ metagene between late fetal and childhood (Figure 2A). Additionally, the loading of the 17^th^ metagene is high at the early fetal and fluctuates around zero at later stages (Figure 2A).

**Figure 2.**
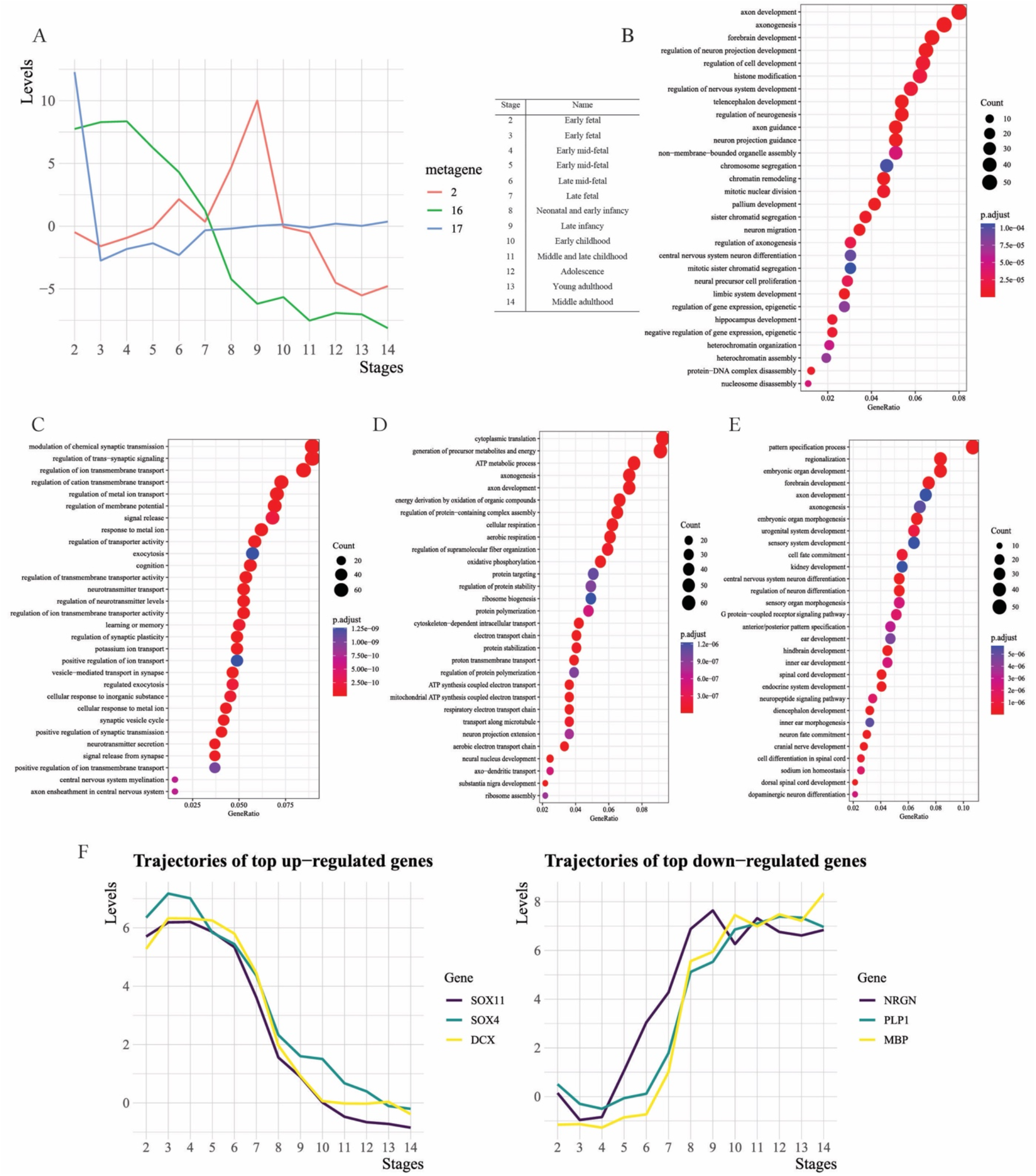
Downstream analyses with results from the application to the BrainSpan data 1. The trajectory of the selected metagene across human brain development are displayed in the left of Figure A. The definition of stages is provided in the table on the right of Figure A. 2. The top 30 up- (Figure B) and down-regulated (Figure C) BPs were enriched by the most variable (16^th^) metagene. 3. The top 30 up- and down-regulated BPs in the M1C-S1C are shown in Figures D and E, respectively. 4. Trajectories of the adjusted expression for the selected genes at the whole brain level are shown in Figure F. The definition of stages in the figure is the same as in Figure A.

To decode biological mechanisms behind these metagenes, we carried out enrichment analyses to find up- (figure 2B) and down-regulated (figure 2C) biological processes (BPs) encoded by the most variable metagene. Here up- or down-regulated BPs refer to BPs enriched by up- or down-regulated genes. Throughout our work, we use them to convey the ideas without loss of clarity. Technical details for the analyses are presented in the section of the enrichment analysis in the Supplementary Content (SC). The up- and down-regulated BPs for the 2^nd^ (Figures S1A and S1B) and 17^th^ (Figures S1C and S1D) metagene were also provided in SC.

On the one hand, a joint analysis of the trajectories of the metagenes (Figure 2A) with their corresponding up-regulated BPs (Figure S1C and 2B) shows that the embryo is under rapid differentiation and development at the early fetal (Figure S1C) and that developments of brain regions, such as the hippocampus, pallium, and forebrain, are high before the mid-fetal period (Figure 2B). As core systems of the brain are established before birth [26], levels of BPs involved in axonogenesis, myelination, and gliogenesis fall rapidly after the late fetal period (Figure 2A and S1C). The basic structure of human brains does not undergo substantial changes after birth, so we observed a decline in levels of BPs involved in the development of brain regions, such as the forebrain, telencephalon, pallium, and hippocampus (Figures 2A and 2B). With its continued development, human brains mature well into the 20s [27], corresponding to a relatively low level of these BPs related to brain development in young adulthood (Figures 2A and 2B).

On the other hand, we observed that the BPs involved in learning, memory, and cognition are relatively low in fetal stages (Figures 2A and 2C, and S1D). With the development of human brains, we also noticed a rapid surge in the level of BPs related to learning, memory, and cognition during infancy (Figures 2A and S1A) and a stable rise in levels of these BPs after mid-fetal stages (Figures 2A and S2C). After the maturation of human brains, these BPs maintain a high level (Figures 2A and 2C). During the development of learning, memory, and cognition, new neural networks form in the brain [28], and the level of neurotransmitters increases simultaneously [29]. Our findings also show that the BPs involved in synaptic functions and neurotransmitter activities follow a similar trajectory as those of learning, memory, and cognition (Figures 2A, 2C, S1A, and S1D).

The trajectory of the least variable metagene is highly positive and flat across human brain development (Figure S2A in SC). Subsequent enrichment analysis revealed that the up-regulated BPs enriched by metagene are involved in cell and ATP metabolism and cell communication (Figures S2B and S2C in SC), which generally maintain a high level during brain development.

### Metagenes characterize development stages and brain regions

Previously, metagenes depict the development trajectories of human brains. Moreover, metagenes also can characterize development stages and brain regions. This can be achieved by examining BPs enriched by metagenes with highly distinct values for development stages or brain regions. We highlighted several metagenes for further study (Tables S1a and S1b in SC).

On the one hand, two metagenes were highlighted to characterize development stages. First, we observed that the loadings of the 5^th^ metage for the early mid-fetal (the 5^th^ stage in Figure 2A) and the 8^th^ metagene for the middle and late childhood are highly positive and much greater than for other stages (Table S1a). Subsequently, the top up-regulated BPs by the 5^th^ metagene involve ossification and development of bone, muscle, and sensory systems (Figure S3A in SC). The results suggest that the mid-fetal period is vital for bone and muscle formation and contributes to sensory system development. Additionally, the 8^th^ metagene encodes BPs up-regulating human metabolic processes (e.g., steroid, fatty acid, cholesterol) and immune responses (e.g., responses to virus and bacteria and leukocyte chemotaxis) (Figure S3B in SC) and down-regulating BPs related to secretion and transport of hormones (e.g., insulin and peptide) (Figure S3C in SC).

On the other hand, we also highlighted three metagenes to uncover the functions of brain regions. The loadings of the 3^rd^ metagene for MD (mediodorsal nucleus) and 12^th^ metagene for CB (cerebellum) are highly negative (Table S1b), but the loading of the 1^st^ metagene for the striatum is highly positive. Furthermore, the absolute values of the three metagenes are much greater for the corresponding brain regions than for other brain regions (Table S1b). As shown by subsequent enrichment analysis, the top 30 down-regulated BPs by the 3^rd^ metagene involve (cardiac) muscle and heart contraction, muscle system process, and regulation of heart rate (Figure S3D in SC), uncovering the key role of MD on motor control.

Additionally, the top 30 down-regulated BPs by the 12^th^ metagene involve synaptic transmission, and sensory and visual system developments (Figure S3E in SC), consistent with the function of CB in receiving inputs from sensory systems and guiding motor control. Moreover, the top 30 up-regulated BPs by the 1^st^ metagene cover learning or memory, cognition, and hormone (e.g., dopamine, monoamine, and catecholamine) secretion and transport (Figure S3F in SC), suggesting that the striatum is responsible for human emotion and cognition (e.g., reward system).

### Reveal trajectories of genes across human brain development at the whole-brain level

Previously, we showed that INSIDER not only enables us to reveal development trajectories with metagenes but also helps to characterize development stages and brain regions. Moreover, INSIDER also allows us to visualize adjusted expression changes of specific genes across human brain development at the whole-brain level.

The development of human brains is heterogenous to different individuals and not linear across different brain regions. Thus, the confounding variation from donors and tissues makes it difficult to reveal gene expression changes at the whole-brain level across human brain development. One can only show trajectories of gene expression in a tissue-wise manner, as displayed in the study [25].

However, INSIDER allows us to obtain the ‘adjusted’ gene expression profiles across development stages by multiplying *D* with *V*. Note that the variation from tissue and donor is controlled for the adjusted gene expression profiles, which cannot be readily computed from conventional approaches. Then, we showed the trajectories of summarized expression changes for six selected genes (Figure 2F), which are chosen from genes in the top and bottom of the expression profile for the most variable (16^th^) metagene selected previously.

The left panel in Figure 2F showed that the trajectory of *DCX*, which is expressed in neuronal progenitor cells and immature migrating neurons, is similar to that of BPs involved in human brain development discussed previously and consistent with its trend in the hippocampus [25]. Regarding *Sox4* and *Sox11*, these are transcription factors known to be key controllers of embryonic and fetal development, and their expression levels are reported to increase simultaneously in the developing central nervous system (CNS) in a previous study [30]. Our results also suggest that they are at high levels in the early development stages (left panel in Figure 2F). Furthermore, they also demonstrate a similar trajectory across human brain development in the panel.

Both *MBP* and *PLP1* are involved in the myelination in the CNS, which is vital to cognition development and function and is largely completed in early adulthood. Another gene *NRGN* regulates synaptic plasticity crucial for synaptic function. The trends of trajectories of the three genes are similar and upward (right panel in Figure 2F). This observation is consistent with the development of cognition discussed previously.

### Uncover biological mechanisms underlying the development of specific brain regions

The tissue representation from INSIDER enables us to explore BPs in specific brain regions by computing their adjusted gene expression profiles. Specifically, the adjusted gene expression profiles for brain regions were obtained by multiplying *T* with *V*. In the operation, two metagenes with the least variance across different brain regions were excluded because they aggregate the expression of cell metabolic and ATP metabolic processes and cell communication and thus excluding them enables us to better uncover BPs related to brain development.

Then, we conducted enrichment analyses on the adjusted gene expression profile of M1C-S1C (primary motor and somatosensory cortex) to explore up- (Figure 2D) and down-regulated (Figure 2E) BPs for the brain region. First, we have no preference for the brain region for further analysis. The main reason is that its function is more specific and less overlapped with other regions, so the validity of our results is more direct.

The M1C-S1C is mainly responsible for generating signals to direct the body’s movement and for processing somatic sensations. The development of this region involves axonogenesis and regulation of neuron projection development (Figure 2D) and thus increases connectivity between synapses. The levels of synaptic activities also increase with the increased connectivity. Therefore, BPs involved in synaptic activities, such as chemical synaptic transmission, trans−synaptic signaling, and synaptic plasticity, are upregulated in this brain region (Figure 2D). However, the levels of BPs involved in the development of human organs, such as the kidney, renal system, spinal cord, and sensory organ (e.g., inner ear and ear), are low in the brain region (Figure 2E).

### Analysis of the aging, dementia, and TBI data

Dementia is a kind of neurodegenerative disorder, and there are various underlying causes. Traumatic brain injury (TBI) may lead to dementia, but the most common cause is Alzheimer’s disease (AD). The Aging, Dementia, and TBI Study aims to characterize neuropathologic, molecular, and transcriptomic changes in the brains of control subjects and TBI cases from an aged population-based cohort [31], which offers RNA-Seq data of temporal cortex (TC), parietal cortex (PC), cortical white matter (WM), and hippocampus (HPC) of controls and TBI cases.

We downloaded the aging, dementia, and TBI data from the website provided in [31]. The data in our analysis contains 377 RNA-seq samples from 107 donors with expression levels of 44,477 genes. In this analysis, we considered the act demented provided for donors as the diagnosis of dementia. We seek to explore BPs contributing to the progression of dementia and discover BPs contributing to dementia in specific brain structures. Here we modeled four covariates: donor, brain structure, phenotype, and gene, and incorporated interactions between brain structures and phenotypes. The number of ranks *K* of latent space selected by hyperparameter tuning is 23.

### Clustering analysis of donor representations

It is challenging to perform clustering analysis in high-dimensional settings, for example with high-dimensional omics data. However, INSIDER enables us to cluster donors with donor representations to characterize donors and even identify disease subtypes, in line with personalized treatment delivery. In practice, we performed hierarchical clustering with latent donor representations (Figure 3A) without the least variable metagene, which encodes BPs related to ATP metabolic processes. We selected two as the number of clusters. Subsequently, we examined the clinical or demographic relevance of the clusters. The two-tailed standard (P=0.0251) and rank-based ANOVA tests (P=0.0185) indicate that the clustering memberships statistically correlate with donor age distribution. Technical details of the statistical tests are provided in SC.

**Figure 3.**
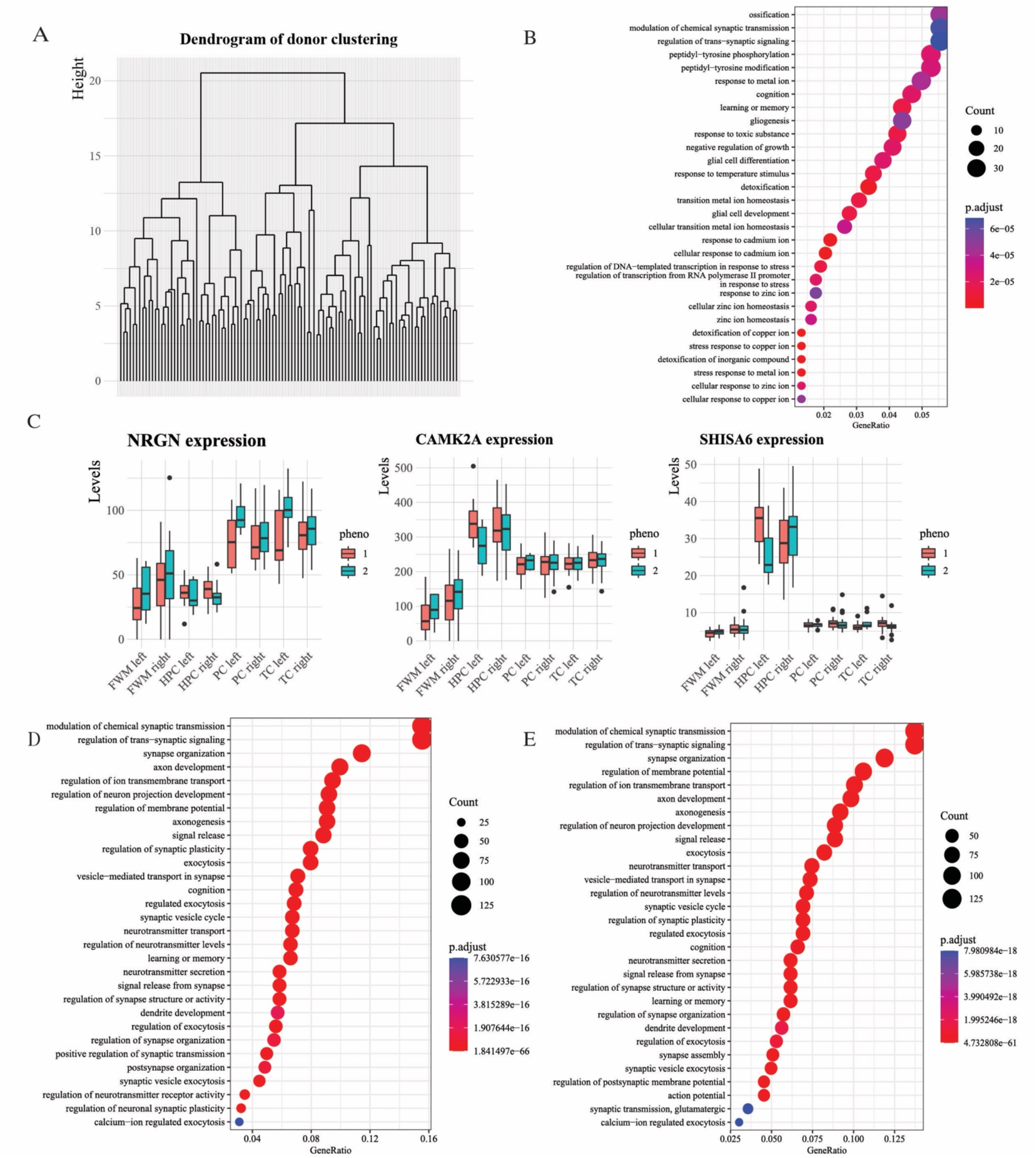
Downstream analyses with results from the application to the aging, dementia, and TBI data 1. Figure A shows the dendrogram of hierarchical clustering with donor representations. 2. Figure B lists the top 30 down-regulated BPs enriched for the difference in gene expression between dementia and control. 3. Figure C compares the expression of three genes between dementia and control across brain regions. Note that “pheno” in the figure is short for “phenotype”. For its values, “1” stands for control and “2” for dementia. “FWM” is short for the white matter of the forebrain, “HPC” for the hippocampus, “TC” for the temporal neocortex, and “PC” for the parietal neocortex. 4. Figure D shows the top 30 down-regulated BPs enriched by the difference in gene expression of left HPC between dementia and control. 5. Figure E shows the top 30 down-regulated BPs enriched for the expression profile for the white matter of the left forebrain.

We also explored whether donor representations enable us to reveal subgroups related to subtypes of dementia. Since not all metagenes are associated with dementia, we filtered off several metagenes that have little relevance to dementia by examining the relevance of the top 30 up- and down-regulated BPs for each metagene. Then, we performed hierarchical clustering on donor representation with these kept metagenes. The dendrogram (Figure S4A in SC) indicates three subgroups.

Two-tailed standard (P=0.0205) and rank-based (P= 0.0731) ANOVA tests yielded similar *p*-values when we examined the association between the clustering and age distribution. In further exploring its relevance to the clinical diagnoses of dementia, Fisher’s exact tests revealed that clusters have a statistically significant association with the Braak staging of dementia (P=0.0025) and are associated with diagnoses of dementia by the NINCDS-ADRDA Alzheimer’s criteria (P=0.108), one of the most widely used criteria in the diagnosis of Alzheimer’s disease. This implies that INSIDER helps to reveal subtypes of dementia.

### Metagenes reveal the biological mechanism of dementia

Here we explored the BPs significantly down-regulated in dementia compared with controls to shed light on the mechanism of dementia. In practice, we selected the top 2 variable metagenes in phenotype representation, as they achieved the most significant p-values in the following enrichment analysis. Then, the adjusted gene expression profiles for dementia and control were obtained by multiplying phenotype representations by gene representation with only the selected metagenes. Subsequently, we examined down-regulated BPs enriched by the difference in gene expression between dementia and control (Figure 2B).

The top 30 BPs enriched for down-regulated genes are statistically significant at the 5e-5 level. We observed that levels of BPs involved in cognition, learning, and memory were significantly lower in the dementia brain than in the control brain (Figure 2B). In addition, synapse activities, such as trans-synaptic signaling, and chemical synaptic transmission, were more severely hindered in the dementia brain than in the control brain (Figure 2B). This observation is supported by previous studies [32]-[34], suggesting that synaptic signaling and transmission play indispensable roles in the nervous system function, and their dysfunctions are known contributing factors to dementia.

Interestingly, expression levels of genes involved in ossification are also significantly different between dementia and control (Figure 2B). The connection between ossification and dementia has been studied in wet lab studies [35], suggesting that multiple risk genes/loci identified in Alzheimer’s disease (AD), such as *APOE, TREM2*, and *CD33*, encode proteins critical for bone homeostasis. Moreover, a degree of comorbidity of AD and osteoporosis is also supported by epidemiological studies [36]. On the other hand, several BPs related to metal ions are also identified in the enrichment analysis (Figure 2B), suggesting that levels of these BPs are lower in dementia than in normal. Since metal ions play essential roles in neurotransmitter synthesis and energy metabolism, their deficiencies contribute to cognitive decline and memory loss [37], which are the most common symptoms of dementia.

### Metagenes characterize donors and functions of brain regions

In addition to revealing the mechanism of dementia, metagenes can help us to characterize donors and reveal functions of brain regions among the aged donors.

We first employed standard two-tailed t-tests to identify metagenes associated with donor characteristics. Statistical test results showed that donors who experienced TBI with loss of consciousness (LOC) have a statistically greater loading in the 13^th^ metagene than those not experiencing (P=0.0242). Subsequent enrichment analysis with the 13^th^ metagene reveals that the top 30 down-regulated BPs involve cognitive functions (Figure S4B in SC), suggesting that experiencing TBI with LOC may be associated with cognitive dysfunction.

Then, we highlighted two metagenes (17^th^ and 18^th^) to reveal the functions of the white matter (WM) and hippocampus (HPC), respectively (Table S2 in SC). We observed that the loadings of the 17^th^ metagene are negative only for WM of both the left and right forebrain, and its value for the WM of the left forebrain roughly doubles that for the right counterpart (Table S2). Similarly, the loading of the 18^th^ metagene is highly positive only for (both left and right) HPC (Table S2).

Subsequent enrichment analysis with the 17^th^ metagene revealed that the top 30 up-regulated BPs involve learning, memory, cognition, and synaptic functions (Figure S4C in SC). This suggests that levels of these BPs are low in both areas of WM than in other brain regions, potentially indicating the involvement of WMs in dementia. Moreover, we noticed that the loading of this metagene is lower in the WM of the left forebrain than in its counterpart. However, this cannot lead us to conclude that WM of the left forebrain is more vulnerable than WM of the right forebrain to dementia, since a single metagene may not show the whole picture of the involvement of WMs in dementia. We will try to clear the issue below.

Besides, the up-regulated BPs by the 18^th^ metagene are related to synaptic functions (e.g., synaptic signaling, transmission, and plasticity), learning, memory, and cognition (Figure S4D in SC), revealing the key functions of the HPC.

### Compare gene expression differences between dementia and control across brain regions

Previously, we explored the overview of dementia throughout all the brain regions studied. Following it, we further examined whether gene expression across different brain regions is similarly affected by dementia. To answer this question, we first selected three genes (*NRGN, CAMK2A*, and *SHISA6*), which were enriched for BPs related to cognition in analyzing biological mechanisms of dementia. Technical details on the gene selection are shown in SC. We plotted the expression levels of the three genes between dementia and control across brain regions (Figure 3C).

Surprisingly, the original expression levels of *CAMK2A* and *SHISA6* in the left HPC were significantly lower in dementia than in controls, but a similar phenomenon was not observed in the right part (Figure 3C). The three genes have been suggested to participate in the progression of dementia or neurodegeneration [38]-[40]. Moreover, the left HPC was postulated to be more vulnerable than the right one to AD [41]. Experimental studies found that the left HPC was affected first by dementia, and an atrophic change in the right HPC was observed after a time lag [42], [43]. In short, our finding seems to suggest that the left and right HPC are affected differently by dementia. We also noticed that no obviously similar phenomenon for *NRGN* was observed (Figure 3C), so we further studied the issue below.

### Discover heterogeneous effects of dementia on brain structures

Previously, dementia was shown to have potentially heterogeneous effects on the left and right HPC at the gene level, so here we seek to compare BPs down-regulated by dementia in the left and right HPC to further clarify the question.

First, we used the left HPC as an example to show how to obtain BPs down-regulated by dementia. First, we multiplied the submatrix *W*_*k*_ of interaction representation corresponding to the left HPC by gene representation *G*. In the operation, several metagenes with the least variance in the submatrix were excluded. Technical details of metagene selection are provided in SC. Then, we examined the down-regulated BPs enriched by the difference in expression profiles of the left HPC between dementia and control (Figure 3D). Similarly, BPs down-regulated by dementia in the right HPC can be obtained (Figure S4E in SC).

Overall, most down-regulated BPs in the left HPC (*P* at 1e-66 in Figure 3D) are more significant than in the right HPC (*P* at 2e-57 in Figure S4E). Moreover, the gene ratio in Figure 3D is also greater than in Figure S4E. Here gene ratio is the proportion of genes enriched for a BP. Regarding BPs enriched for the two brain structures, there is little difference in general. This leads to a similar finding that the left and right HPC are affected differently by dementia and supported by previous studies we discussed earlier.

### Uncover BPs in specific structures of the aged brain

Besides, INSIDER also enables us to find BPs up- and down-regulated for a specific region. Here we are interested in key BPs in the white matter (WM) among the aged, as white matter plays an essential role in learning and brain functions. Our previous analysis shows that left hemisphere involvement is more likely to result in disabling cognitive functions in dementia, so we seek to explore whether a similar phenomenon can be observed in WMs, an issue we raised previously. We examined BP down-regulated in WM of both the left and right forebrain.

First, we computed the adjusted gene expression profiles for the WM of both the left and right forebrain by multiplying the brain structure representation with the gene representation. In the multiplication, only the top 3 variable metagenes in terms of brain region representations were included. Then, we conducted enrichment analyses on the expression profile of WMs of both the left and right forebrain to reveal the top 30 down-regulated BPs (Figures S4F in SC and 3E).

The BP down-regulated in WM of the left forebrain (Figure 3E) is slightly more significant than that in the right counterpart (Figure S4F). Moreover, the BPs down-regulated in both areas are basically the same. Our analysis may suggest that both areas of the WM are similarly affected by dementia.

### Analysis of gender differences in brain expression with GTEx

In this application, we seek to explore gender differences in gene expression across 13 different brain regions from the GTEx data [2]. Details of data processing are shown in SC. The processed data contains 988 expression profiles of 49,999 genes from 318 different donors. For each brain region, it has 38 RNA-seq samples from either men or women. We modeled three covariates: brain region, gender, and gene. The number of ranks *K* of latent space selected by hyperparameter tuning is 12.

### Metagenes reveal gender differences in brain expression

To reveal gender differences in brain expression, we calculated the adjusted gene expression profiles for genders by multiplying gender representation with gene representation and then examined up- (Figure 4A) and down-regulated (Figure 4B) BPs enriched by the difference in gene expression between the male and the female.

**Figure 4.**
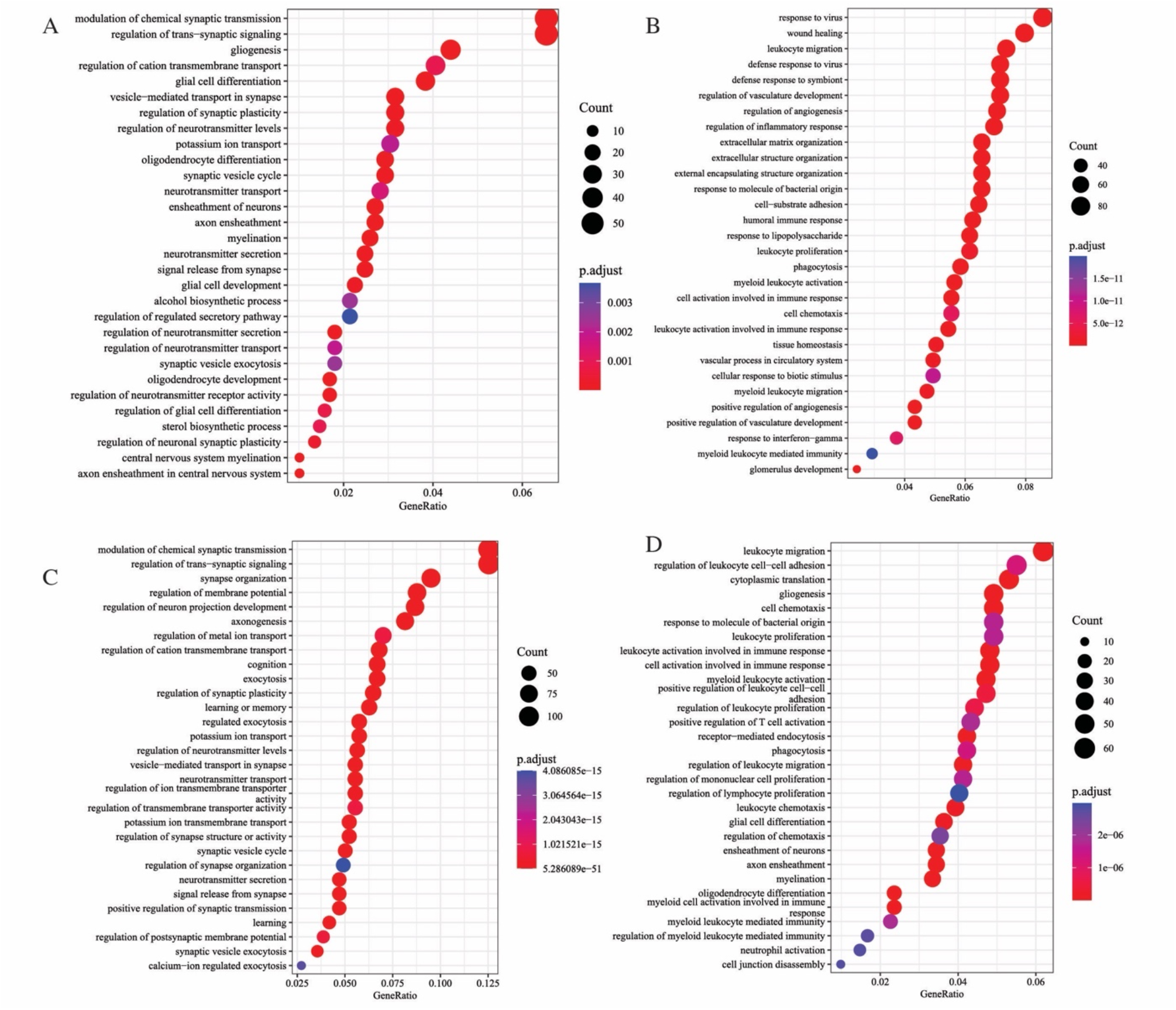
Downstream analyses with results from the application to the GTEx data 1. Figures 4A and 4B show the top 30 up- and down-regulated BPs enriched by the difference in gene expression between genders, respectively. 2. Figures 4C and 4D display the top 30 up- and down-regulated BPs enriched for the difference in gene expression of the frontal cortex (BA9) between genders, respectively.

On the one hand, expression levels of genes enriched for synapse activities and neurotransmitters are higher in men than in women across the brain regions included (Figure 4A). On the other hand, women have higher expression levels of genes contributing to immune responses, such as (defense) response to viruses and bacterium and humoral immune response, than men (Figure 4B).

### Metagenes characterize functions of brain regions

Previously, we revealed gender differences in brain expression utilizing metagenes. Additionally, metagenes also enable us to uncover the functions of brain regions included. Intuitively, if the loading of a specific metagene for one brain region is significantly greater than that of other brain regions, then the metagene can be utilized to characterize the brain region. Following the intuition, we highlighted several metagenes (Table S3 in SC) to demonstrate how to achieve this goal in practice.

First, we found that the loading of the 3^rd^ metagene is highly positive and much greater in the three subcortical nuclei (putamen, nucleus accumbens, and caudate) than in other brain regions (Table S3). The top 30 BPs enriched by up-regulated genes of the metagene mainly fall into the following aspects: muscle contraction and system process, blood circulation and circulatory system, response to monoamine, catecholamine, and dopamine, synaptic transmission and signaling, and neurotransmitter transport (Figure S5A in SC), highlighting their roles in human motor and emotions.

Meanwhile, we also examined the top 30 up-regulated BPs enriched by the 4^th^ (Figure S5B in SC) and 6^th^ (Figure S5C in SC) metagenes since the loadings of the 4^th^ metagene for the hypothalamus and 6^th^ metagene for HPC are highly positive and much greater than for other brain regions (Table S3). The up-regulated BPs enriched by the 4^th^ metagene primarily involve hormone secretion and transport, calcium ion homeostasis, and catecholamine and monoamine transport (Figure S5B), which is consistent with the role of the hypothalamus in homeostasis [48]. Meanwhile, the up-regulated BPs enriched by the 6^th^ metagene involve learning and memory, synaptic transmission and signaling, and dendrite development (Figure S5C), revealing the key functions of HPC.

Moreover, we also explored the top 30 up-regulated BPs enriched by the 12^th^ metagene, whose loading is highly negative in the spinal cord. The up-regulated BPs involve the following biological activities: learning, memory, cognition, synaptic transmission and signaling, and neurotransmitter transport and secretion (Figure S5D in SC), implying a much lower involvement of the spinal cord in human cognitive function, compared with other brain regions.

### Explore gender differences in the expression of specific brain regions

In the previous subsection, we revealed gender differences in brain expression across all brain regions included. Still, psychiatrists might be more interested in the gender difference in a specific brain region since genders show differences in mental disorder prevalence [49]. Uncovering gender differences in the expression of a particular brain may help us understand gender differences in mental disorders.

First, we identified the brain structure that shows the greatest difference between genders. Specifically, for a specific brain structure *i*, we calculated the Euclidean distance between the representations of the structure for male *w*_*i*1_ and female *w*_*i*2_ with the interaction representation *W*. Then, we selected the brain structure showing the greatest distance. The frontal cortex (BA9) showed the greatest distance and was chosen for further investigation.

After choosing the frontal cortex, we computed its adjusted gene expression profiles for both males and females by multiplying the submatrix of *W* corresponding to the frontal cortex by the gene representation *V* only on selected metagenes. In determining metagenes, we considered the top 2 metagenes showing the greatest difference between genders using the submatrix corresponding to the frontal cortex. Then, we examined the up- (Figure 4C) and down-regulated (Figure 4D) BPs enriched by the difference in the adjusted gene expression of the frontal cortex between genders.

Generally, the main theme of up- and down-regulated BPs (Figures 4C and 4D) for gender differences in expression of the frontal cortex are similar to those in Figures A and B. Further results regarding gender differences in BPs in BA9 can be found in the two figures. Note that the BA9 area is only one part of the frontal cortex, so the results may be biased and incomplete in revealing the whole picture of the frontal cortex.

## Discussion

We demonstrated the versatility and potential of INSIDER through three applications, in which various downstream analyses were performed. In the first application, the development trajectories across the human life span were revealed, and up- and down-regulated BPs underlying them were uncovered. In two other applications, we explored the mechanisms contributing to dementia and gender differences in brain expression, identified subgroups among donors with latent donor representations, and examined the contribution of tissues (e.g., brain regions) to biological conditions (e.g., dementia and gender). Our results are supported by previous studies. For example, the interpretation of the development trajectories is consistent with findings from previous studies.

INSIDER has the following advantages. First, it is general and flexible, in which covariates and interactions in modeling can be adjusted according to research interest. Second, it is versatile and interpretable in terms of downstream analysis. Various downstream analyses were carried out, such as revealing development trajectory, clustering donors, revealing disease mechanisms, and exploring interactions between covariates. Third, it encourages sparsity in gene representations by introducing the elastic net regularization, thus facilitating result interpretations. Fourth, it can capture interactions between covariates, common in biological settings. Finally, it has good scalability and allows missing values.

Bulk RNA-Seq techniques are popular in studying tissue-specific expression profiles. Usually, confounding variables (covariates) are inevitably introduced in the RNA-Seq expression data. For example, GTEx [2] data contains confounding variables, such as donor, gender, and age, which significantly impact tissue expression, as demonstrated in our study. However, few studies work on the issue. Existing approaches such as PCA, factor analysis, and non-negative matrix factorization only can reduce data dimensionality but cannot tackle it. It has become more prominent with the increasing availability of RNA-seq data and the popularity of joint data analysis from different sources. For example, the PsychENCODE project [4] is composited of several independent projects, such as ATP (the Autism Tissue Program) [5], CMC (the CommonMind Consortium) [51], and BrainGVEX [4], with different study aims. INSIDER well addresses the challenge and takes account of sparsity in gene expression and the interaction between covariates simultaneously. More importantly, it has excellent interpretability and can be applied to large-scale data analysis. Future works may include further analysis of various bulk RNA-seq datasets and extend it to single-cell data analysis.

## Conclusion

This article proposed a novel statistical approach, INSIDER, based on matrix decomposition, to model the variation contributed by confounding variables or covariates in bulk RNA-Seq data and decompose it into a shared low-rank latent space. The variation may originate differently, such as donor, gender, tissue, or other biological conditions. INSIDER is general and flexible framework, which is robust to missing values. In particular, it can capture interactions between variables or covariates. Moreover, it encourages sparsity in latent representations by imposing the elastic net penalty, thus facilitating model interpretations. We employed it to three different RNA-Seq data, and various downstream analyses were carried out based on results from INSIDER. The broad applicability of INSIDER in biomedical and clinical settings was demonstrated in the applications.

## Methods

Here we propose INSIDER, which uses additive matrix factorization to model heterogeneous biological variation in RNA-Seq data. We assume that heterogeneous variation shares a low-rank latent space since the heterogeneous variations are usually contributed by changes in the same signaling pathways across individuals with different phenotypes. A study [52] also showed that the low-rank latent representations, also known as metagenes, aggregate the information of gene expression and can help gain insights into the biological mechanisms of interest.

### Model Specifications

Denote *z*_*ijhm*_ as the expression level of gene *m* in tissue *h* of donor *i* with phenotype *j*. We assume that there are *N*_1_ donors, *N*_2_ phenotypes, and *N*_3_ tissues. *z*_*ijhm*_ is modelled as

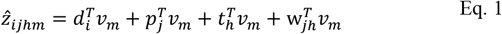

where *d*_*i*_, *p*_*j*_, *t*_*h*_, *w*_*jh*_, *v*_*m*_ are vectors of length *K*. Note that the above equation incorporates the interaction effect between tissues and phenotypes via introducing *w*_*jh*_, which is mediated by genes, since the interaction may affect different genes differently. Note INSIDER is flexible, and more covariates can be incorporated depending on the study: for example, development stages and treatment assignments can be included as covariates. Here we do not consider the interaction between individuals and tissues or phenotypes, even though more interaction terms can be easily incorporated into INSIDER.

The objective function that we seek to minimize for Eq. 1 is defined as

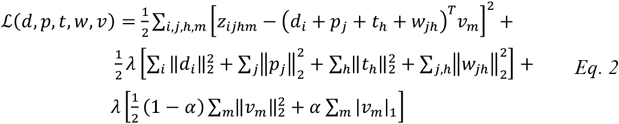

In the above equation, we incorporate 1/2 for easier implementation. The Eq. 2 can be rewritten with matrix notations. Let *Z* ∈ ℝ^*N*×*M*^ represent the data matrix for the expression profile of *N* samples and *M* genes. The matrices 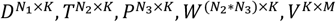 are the rank *K* latent representations for *N*_1_ donors, *N*_2_ phenotypes, *N*_3_ tissues, *N*_2_ ∗ *N*_3_ interactions, and *M* genes, respectively. 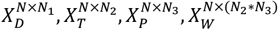 are indicator matrices, which represent the dummy design matrices for *N* samples. Eq. 2 can be rewritten as follows:

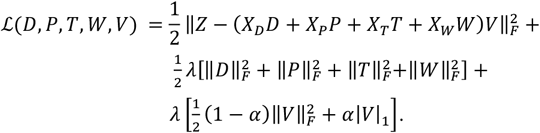

In the above equation, missing entries in *Z* are removed in the summation in the first line in the above equation.

### Model fitting

Alternating block coordinate descent (BCD) was employed to optimize Eq. 2: each time we update a subset of the parameters with all the other parameters fixed. For example, when all parameters except *v*_*m*_ are fixed, then our problem becomes a set of linear regression problems with elastic net regularization and all *v*_*m*_ can be updated simultaneously. In each iteration of BCD, we update all parameters in our model sequentially. We repeat the process until the stop criteria meets, which is presented in the next subsection. The following are the details for minimizing the objective function defined by Eq. 2. using alternating BCD.

### Optimize the objective function

Given the objective function defined by Eq. 2, we choose to maximize the objective function with respect to *p*_*j*_ first. By fixing all *d*_*i*_, *t*_*h*_, *w*_*jh*_, *v*_*m*_ and taking the derivative with respect to *p*_*jl*_, the *l*-th element of vector *p*_*j*_, we have

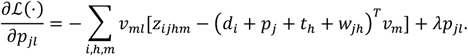

We set the above equation to zero and have

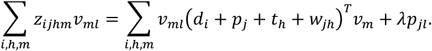

After a simple vectorization of the above equation, we have

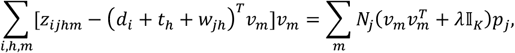

where *N*_*j*_ is the number of samples from donors with phenotype *j*, and 𝕀_*K*_ is an identity matrix of rank *K*. The update for *p*_*j*_ obtained from the above equation is as the following

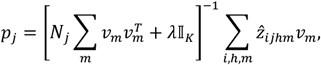

where 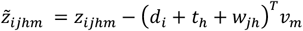. Similarly, the update for *t*_*h*_ can be derived as

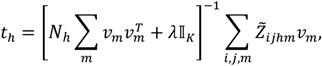

where 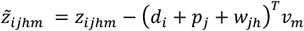 and *N*_*h*_ is the number of samples from tissue *h*. Likewise, we also can easily derive the update for *d*_*i*_ as following:

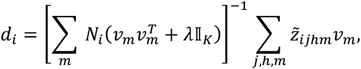

where 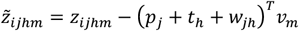, and *N*_*i*_ denote the number of samples from donor *i*. Finally, the update for *w*_*jh*_ can be derived as the following

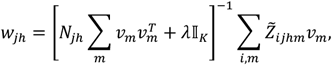

where 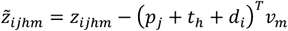 and *N*_*jh*_ is the number of samples of tissue *h* with phenotype *j*.

When we seek to minimize the objective function with respect to *v*_*m*_ with all the other parameters fixed, the optimization problem degenerates to a set of elastic net problems with the same parameterization as that used in the R packages **glmnet** [53]. Following the matrix representation of Eq. 2, the objective function with respect to *V* is

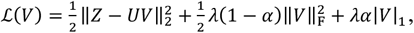

where *U* = *X*_*D*_*D* + *X*_*P*_*P* + *X*_*T*_*T* + *X*_*W*_*W*. Note that *V* is a *K* × *M* matrix, so the above equation defines a set of *M* elastic net problems that correspond to the *M* genes. All the elastic net problems defined by the equation are independent and can be optimized in parallel. The elastic net problem defined by parameters *v*_*m*_, the *m*-th column of *V*, is of the form

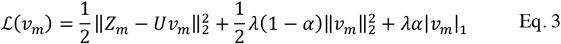

In the equation, *Z*_*m*_ is the expression levels of the *m*-th gene across the *N* expression profiles. Missing entries in *Z*_*m*_ are removed in the above calculation. The randomized coordinate descent (RCD) algorithm proposed in Algorithm 1 is employed to solve the problem defined by Eq. 3.

In practice, we find that optimizing the subproblems with elastic net regularization is computationally intensive. The safe rules for screening are adopted to accelerate the computation, which discard variables with coefficients that are shrunk to zero in optimization. For illustration, we consider the elastic net problem defined by Eq. 3 with *λα* and *λ*(1 − *α*) for L1 and L2 penalties, which is of the same form that was adopted in the **glmnet** package [53]. The strong screening rules defined by Eq. 24 in the study [54] can be directly applied to Eq. 3, and the rule for discarding predictor *j* is

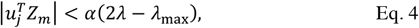

where 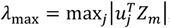 and *u*_*j*_ is the *j*-th column of *U*.

### Initialization, hyperparameter tuning and the stopping criterion

Regarding initialization of the latent representations {*D, P, T, W, V*} defined in the matrix representation of Eq.2, all latent representations are initiated from normal distribution *N*(0, 0.001). For the initialization as a warm start for the subproblem defined in Eq. 3 in optimization, we consider the solution from ridge regression or the solution from the previous iteration, depending on which one leads to a lower loss function defined by Eq. 3. The warm start with the ridge solution for the problem defined in Eq. 3 is as follows

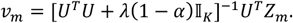

In model selection, grid search is used to select the hyperparameters {*K, λ, α*} for INSIDER.

#### Algorithm 1

RCD algorithm for elastic net with screening rules

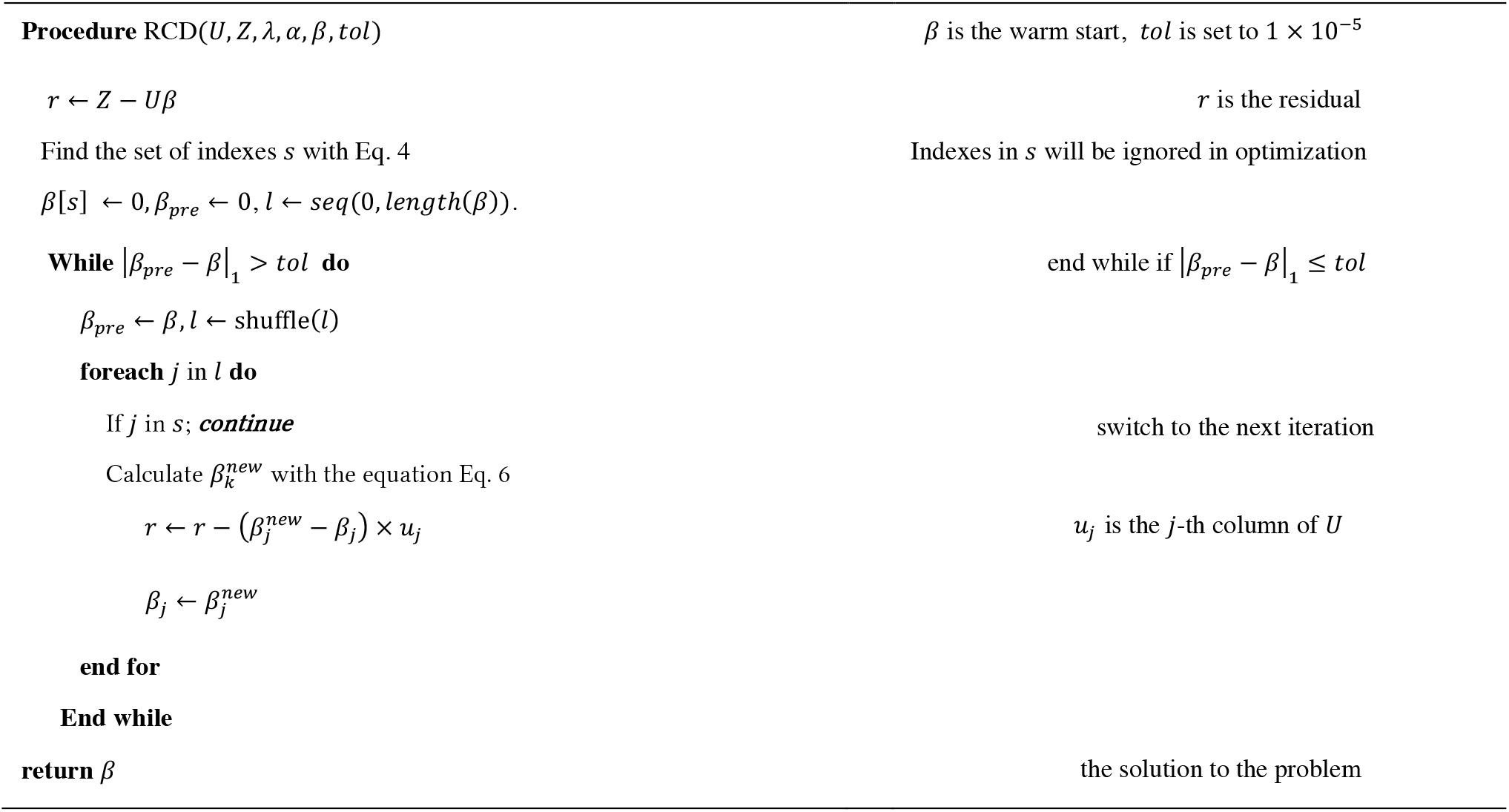

We randomly hold out 10% of the whole dataset as a test set and choose the set of parameters that performs the best on the test set in terms of root-mean-square error (RMSE). For each set of candidate hyperparameters, we run alternating BCD for 50 iterations. In practice, we select *K*, the rank of latent representations, from a sequence of integers from 10 to 30 with step size 1. In the process, *λ* is set to 0.1 to avoid singularity in the matrix inverse, and *α* is fixed to 0. After choosing *K*, we define a broad parameter grid for *λ* (0.1 to 100) and *α* (0 to 1 with step size 0.1) and choose the parameters with the best performance on the test set.

In our study, the relative difference in the objective function between two successive iterations is computed for each iteration, and the stopping criteria using this measurement is defined as

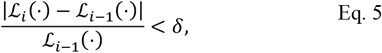

where ℒ_*i*_(⋅) denotes the loss at the *i*-th iteration, and *δ* is a predefined threshold and set to 10^−11^ in our experiment.

## Supporting information

Supplementary Content

## Appendices

### Random coordinate descent for elastic net problems

We define the objective function for the elastic net problem as

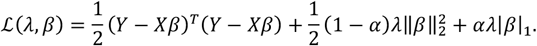

Here *Y* is the outcome, *X* is a *n* × *p* feature matrix, *β* is the parameter vector, and *α* and *λ* are regularization parameters.

Here we derive the update for a single parameter *β*_*j*_ for demonstration. Denote *β*^*k*^ the parameter *β* at the *k*-th iteration. We rewrite the above equation regarding *β*_*j*_ for *k +* 1 iteration as

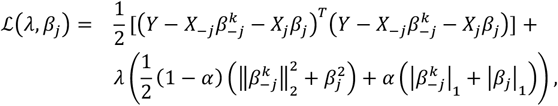

where *X*__ *j*_ is *X* without the *j*-th column, and 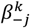 is the *β*^*k*^ with the *j*-th element removed and is known at the *k +* 1 iteration when updating *β*_*j*_ Taking the derivative of the above equation regarding *β*_*j*_ and setting it to zero, we have

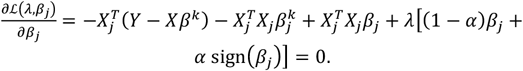

The update for *β*_*j*_ at *k +* 1 iteration is as follows:

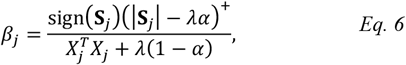

where 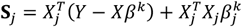

## Acknowledgment

This research has been supported by the Chinese University of Hong Kong startup grant (4930181), the Chinese University of Hong Kong’s Project Impact Enhancement Fund (PIEF) and Science Faculty’s Collaborative Research Impact Matching Scheme (CRIMS), and Hong Kong Research Grant Council (ECS 24301419, GRF 14301120).

