## Supplementary Content for "INSIDER: Interpretable Sparse Matrix Decomposition for Bulk RNA Expression Data Analysis": Supplementary content(SC).docx

Enrichment analysis

Here we focus on the biological processes (BPs) enriched by genes with large effects. Thus, genes from the top or low quantile are used to identify the most impactful BPs encoded by the adjusted expression profile or difference in expression profiles. In the study, the enrichment tests were conducted on the set of genes in the upper or lower 2.5% quantile of the adjusted expression profile or difference in expression profiles to explore the up-regulated or down-regulated BPs, respectively. Even though the threshold for selecting genes for enrichment analysis is somewhat arbitrary, important BPs encoded can be revealed to help us gain insights into the biological mechanisms of our interests. In the implementation, the function *enrichGO* from the R package clusterProfiler (v4.4.4) was utilized to conduct the analysis, with the “ont” parameter equal to "BP" to identify BPs enriched by gene set.

Clustering analysis with donor representations

After performing clustering analysis, we are interested in the clinical or demographic relevance of the clustering. Thus, we performed the ANOVA test to examine the association between the clustering and the age of donors. As there are age ranges available for several donors, the median of age range was considered as the age in conducting the analysis. In practice, the standard and rank-based two-tailed ANOVA tests were carried out. Their results showed that our clustering has a statistically significant association with donor age distribution, with p-values equal to 0.0251 in the standard ANOVA test and 0.0185 in the rank-based ANOVA test. This suggests a statistically significant difference in the age distribution of donors among the two clusters.

Details for gene selection in comparing expression differences between dementia and control across brain regions

As we mentioned in the main text, we first filtered out five biological processes (BPs) related to cognition from the analysis of the biological mechanism of dementia. The five BPs are learning or memory, cognition, response to metal ions, modulation of chemical synaptic transmission, and regulation of trans−synaptic signaling. Then, we collected all genes (~82 genes) enriched for the five BPs. Here we are interested in the genes whose expression are statistically different across brain region. Thus, we conducted t-tests to check the significance of the difference in expression across different brain regions for each gene and then employed Simes’ Test [2] to obtain aggregated p-values for each gene, which controls the Type I error rate by taking account of the issue of multiple testing. Thus, we focus on the genes with aggregated p-values significant at ~0.05. There are five genes left. Finally, three genes (NRGN, CAMK2A, and SHISA6) are selected for demonstration.

### Detail for metagene selection for discovering heterogeneous effects of dementia on brain structures

In practice, we used the following strategy to select metagene selection for the study purpose. First, we obtained the profiles of the left HPC $E$ for dementia and control by multiplying the submatrix $W_{k}$ of interaction representation corresponding to the left HPC by gene representation $G$. In the above multiplication, only top-*N* metagene with the greatest difference in $W_{k}$ were used. The number of genes (*N*) is determined by the significance of p-values in the following-up enrichment analysis. We selected the number of top genes which leads to the most significant p-values. Here, $E$ is a matrix of two rows corresponding to the adjusted expression profiles of the left HPC for dementia and control. Then, we calculated the difference between the two rows of $E$ by $E_{1}-E_{2}$ and examined the down-regulated BPs enriched by the difference. For the right HPC, we follow the exact same procedure as above to ensure consistency.

Data processing for GTEx

We downloaded the data file named “GTEx_Analysis_2017-06-05_v8_RNASeQCv1.1.9_gene_tpm.gct.gz” from the official website (<https://gtexportal.org/home/datasets>). Then, we filtered out the brain expression profiles from the GTEx data for further processing and down-sampled the number of male donors to maintain an even number of donors in both genders across different brain regions. This practice is to ensure that the latent representation of brain regions captures roughly even information regarding brain expression from both males and females.

Figure S1 The top 30 up- and down-regulated BPs enriched by the 2nd and 17th metages

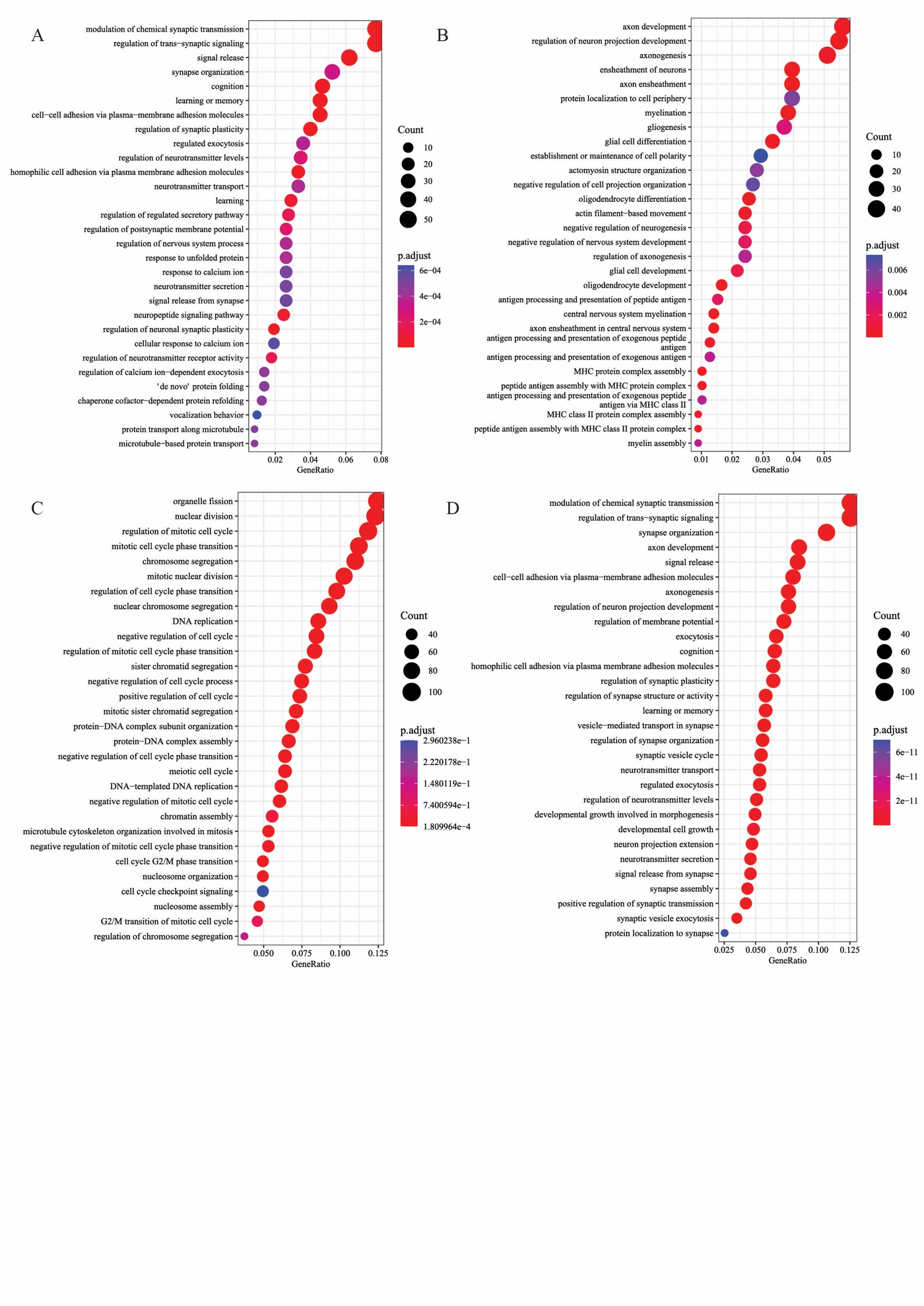

1. The top 30 up- (Figure A) and down-regulated (Figure B) BPs are enriched by the 2nd metagene.
2. The top 30 up- (Figure C) and down-regulated (Figure D) BPs are enriched by the 17th metagene.

Figure S2 The trajectory of the least variable metagene and top 30 BPs enriched by it

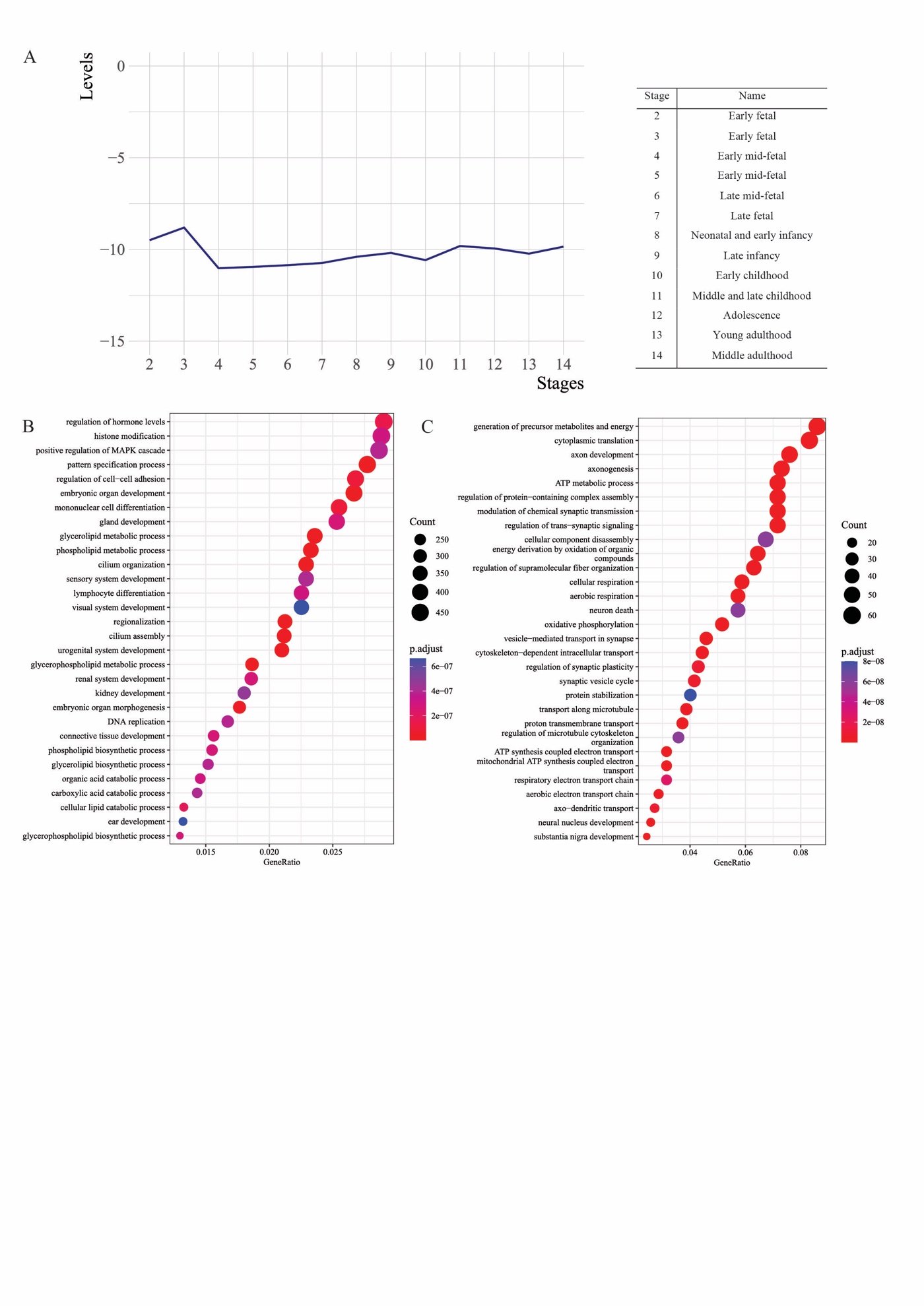

1. Figure A shows the trajectory of the least variable metagene across human brain development.
2. The top 30 up- (Figure B) and down-regulated (Figure C) BPs are enriched by the least variable metagene.

Figure S3 Metagenes characterize development stages and brain regions

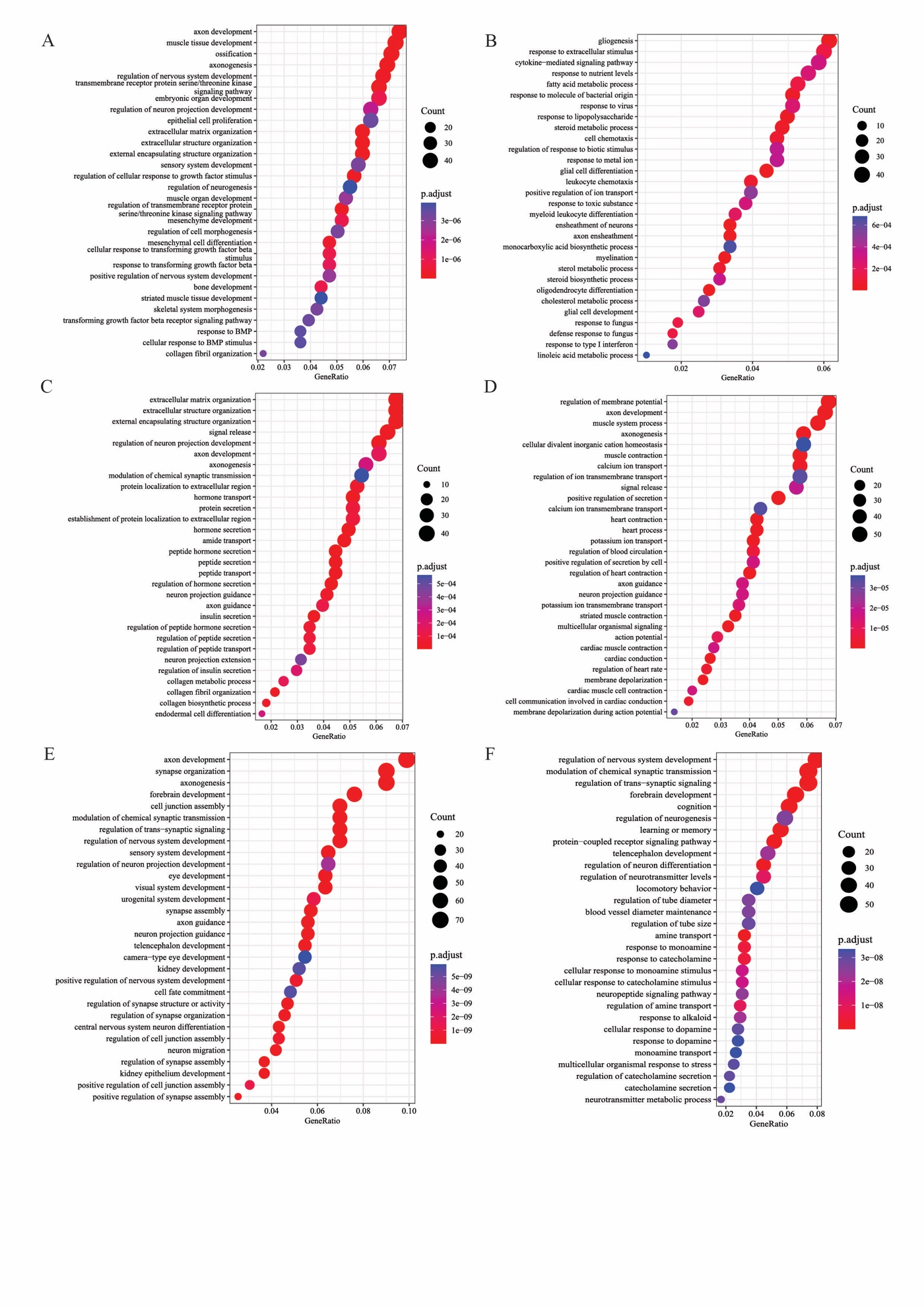

1. Figure A shows the top 30 up-regulated BPs enriched by the 5th metagene.
2. The top 30 up- (Figure B) and down-regulated (Figure C) BPs are enriched by the 8th metagene.
3. The top 30 down-regulated BPs enriched by the 3rd and 12th metagenes are shown in Figures D and E, respectively.
4. The top 30 up-regulated (Figure F) BPs are enriched by the 1st metagene.

Figure S4 Supplement figures for analysis of the aging, dementia, and TBI data

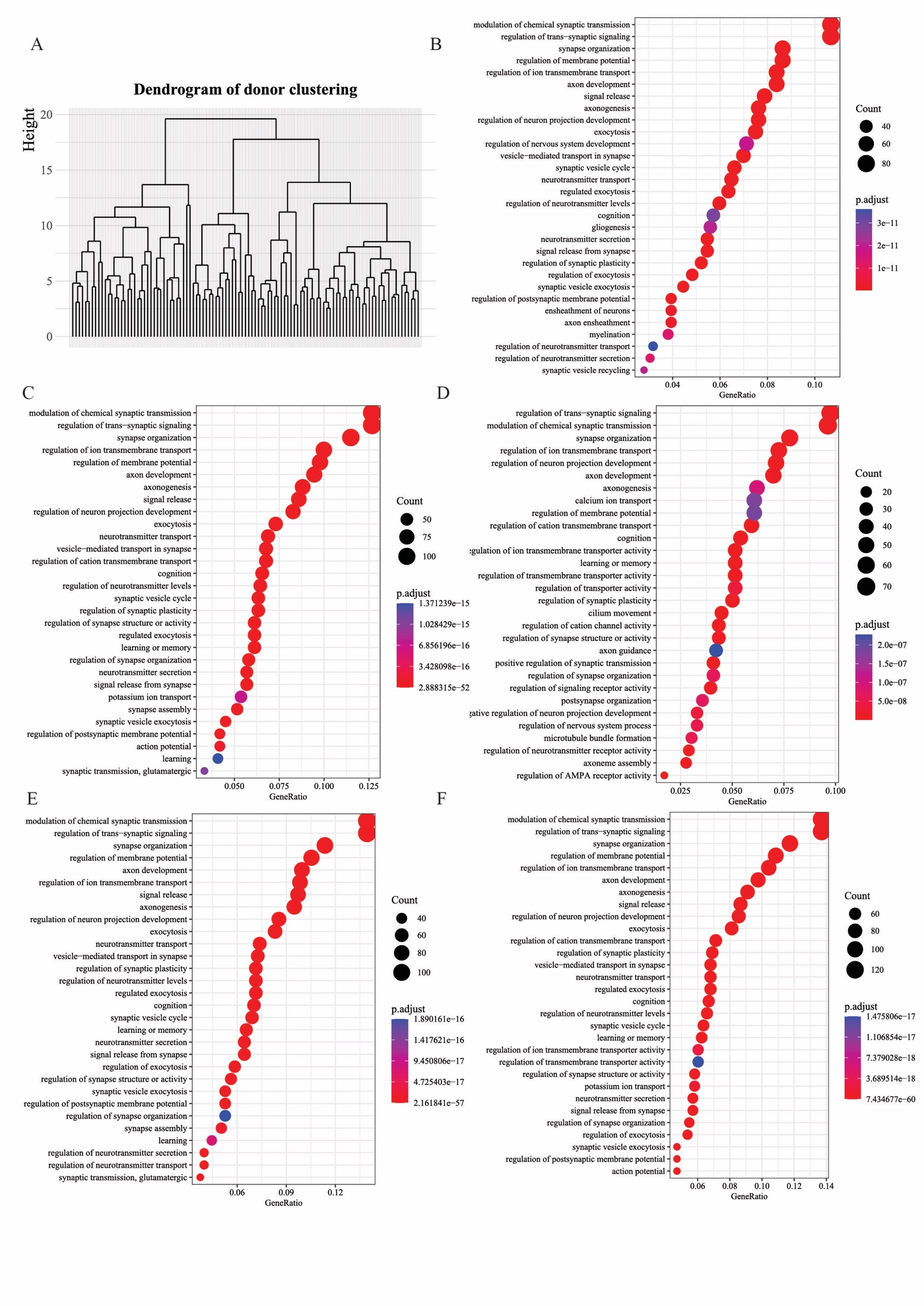

1. Figure A shows the dendrogram of hierarchical clustering on donor representation with the selected metagenes.
2. The top 30 down-regulated BPs enriched by the 13th metagene are shown in Figure B.
3. Figures C and D show the top 30 up-regulated BPs enriched by the 17th and 18^th^ metagenes, respectively.
4. Figures E and F show the top 30 down-regulated BPs enriched by the difference in expression profiles of the right HPC and WM of the right forebrain, respectively.

Figure S5 Supplement figures for analysis of GTEx

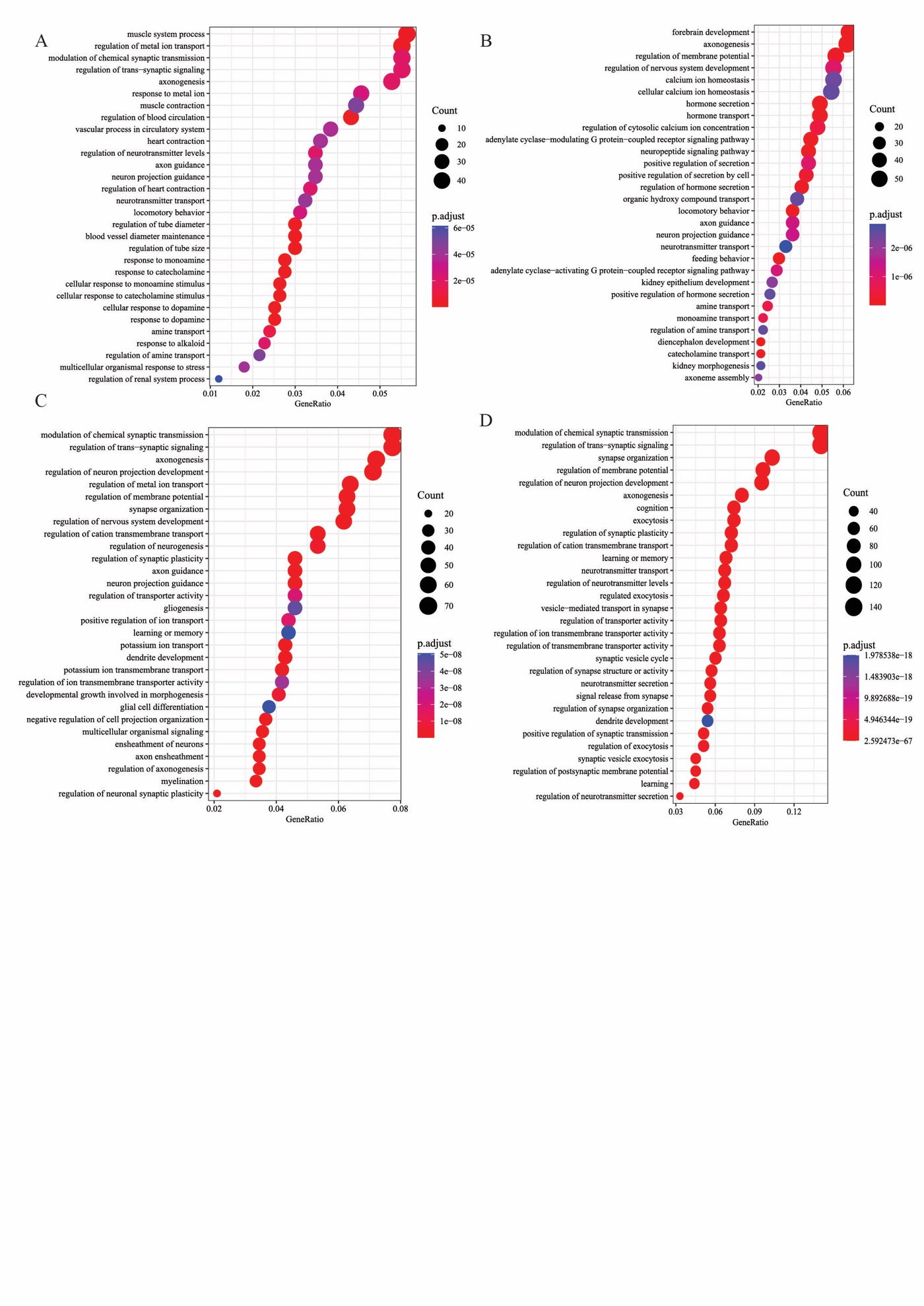

1. Figures A, B, C, and D show the top 30 up-regulated BPs enriched by the 3rd, 4th, 6th, and 12th metagenes, respectively.

stylefix
